# Deep Brain Stimulation Microelectrodes as a Source of Human Subcortical RNA: Validation of a Low-Input Transcriptomic Protocol

**DOI:** 10.64898/2026.07.27.740887

**Authors:** Sandeep Sandeep, Arti Saini, Vivek Chouhan, Sanskriti Rai, Divya Madathiparambil Radhakrishnan, Umran Yaman, Arunmozhimaran Elavarasi, Divyani Garg, Animesh Das, Thyagarajan Radhakrishnan, Anu Gupta, Mamta Bhushan Singh, Venugopalan Yamuna Vishnu, Saroj Kumar, Abhishek Yadav, Ajay Garg, Rohit Bhatia, Ahmadulla Shariff, Achal Kumar Srivastava, Lata Rani, Dattaraj Savarakar, Rajinder Kumar, Manmohan Singh, Kanwaljeet Garg, Kailash Bhatia, Henry Houlden, Poodipedi Sharat Chandra, Roopa Rajan

## Abstract

**Background:** Understanding the molecular basis of Parkinson’s disease (PD) phenotypic heterogeneity maybe improved by in vivo access to deep brain tissue. Deep brain stimulation (DBS) surgery offers a unique opportunity: microelectrodes traversing the subthalamic nucleus (STN) carry adherent brain tissue upon withdrawal, providing a source of RNA from subcortical regions in living patients without additional invasive procedures.

**Aim:** To develop and validate a low-input RNA extraction and transcriptomic profiling protocol using DBS microelectrodes in post-mortem human brain, to confirm that recovered RNA is of brain rather than blood origin, and to characterise its sub-regional and cellular identity.

**Methodology:** DBS microelectrodes were inserted without image guidance or guide tube into three unfixed post-mortem human brains targeting the STN trajectory. In clinical practice, a guide tube shields the electrode from cortical tissue; its absence here means tissue from the full insertion trajectory may contribute to recovered RNA. Thirty-eight microelectrodes were evaluated across single and pooled strategies. RNA was extracted using a modified RNeasy Micro low-input protocol; libraries prepared using NEBNext Single Cell/Low Input RNA Library Kit and sequenced on Illumina NovaSeq 6000 (paired-end, 2×150 bp). Tissue identity was validated against GTEx v10 (54 tissues) and Allen Human Brain Atlas (ABA, 19 subcortical regions including STN) using Pearson correlation with permutation testing (1,000 permutations) and BH-FDR correction. Transcriptional overlap between subcortical and cortical reference regions was quantified and subcortical-enriched gene filtering performed.

**Results:** Twenty-five RNA isolates were obtained from 38 microelectrodes; 54.5% of Bioanalyzer-assessed samples achieved RIN ≥5 (median 7.1; range: 5.9-7.8). Fifteen libraries passed sequencing QC (mean depth 43.0 ± 14.1 million read pairs; mean Q30 83.2 ± 5.6 %). PCA of rlog transformed expression data resolved samples by donor identity. All samples confirmed brain tissue origin by GTEx τ-index tissue specificity correlation (n=996 brain and blood specific markers). Brain correlations were highest for frontal cortex (mean r= 0.683 ± 0.072), anterior cingulate cortex (mean r= 0.658 ± 0.069), amygdala (mean r= 0.615 ± 0.068), and basal ganglia including caudate (mean r= 0.553 ± 0.059), putamen (mean r= 0.546 ± 0.059), and nucleus accumbens (mean r= 0.544 ± 0.057); all BH-FDR < 0.05. Cerebellum showed the lowest brain correlations (cerebellar hemisphere: mean r = 0.327 ± 0.039; cerebellum: mean r = 0.314 ± 0.037). Whole blood correlation was strongly negative (mean r= −0.250 ± 0.079), confirming non-haematological origin. ABA genome-wide analysis confirmed positive STN correlation (r=0.54–0.62; permutation p<0.001). MuSiC cell-type deconvolution against the Allen Brain Atlas HMBA-BG snRNA-seq reference identified oligodendrocytes (33.5 ± 17.2%), frontal cortical neurons (9.1 ± 7.0%), STR D1 MSNs (6.0 ± 4.9%), and dopaminergic neurons (1.1± 2.7%) as the principal cell types, independently corroborating bulk transcriptomic findings at single-cell resolution.

**Conclusion:** This study validates a low-input RNA extraction protocol for DBS microelectrodes, confirming brain-specific transcriptomic profiles consistent with STN-adjacent subcortical sampling. The 96% transcriptional overlap between cortical and subcortical regions, combined with absence of a guide tube in this post-mortem model, limits sub-regional specificity; clinical application with a guide tube would enrich the subcortical signal. These findings provide the methodological framework for in vivo molecular profiling of the human basal ganglia during DBS surgery in PD patients.

## Introduction

Parkinson’s disease (PD) is the second most common neurodegenerative disorder among older adults. Although termed a single disease, PD is enigmatically heterogenous clinically, with presentations ranging from tremor dominant, akinetic rigid and non-motor predominant, among others. (1) The rate of progression to advanced disease also varies. (2) Moreover, the response to medications and deep brain stimulation (DBS), which is the standard of care for patients with disabling motor complications despite best medical therapy, differs. (3) Such heterogeneity is partly explained by genetic variability-several monogenic or risk-associated genes result in distinct clinical phenotypes. (4) *LRRK2* or *PRKN*-associated PD generally respond well to DBS while PD associated with variants in *GBA* or *SNCA* result in parkinsonism with earlier cognitive impairment or autonomic dysfunction and poor response to DBS. (5) However, monogenic causes of PD are infrequently encountered in clinical practice (<5-10%) and the cause for phenotypic heterogeneity in the large, apparently sporadic PD population remains poorly understood. (6)

To understand the brain level mechanisms underlying PD phenotypes, a standard approach is to study tissue morphology and gene expression in post-mortem brains. (7), (8), (9) Early studies on post-mortem brains identified that the pathophysiological hallmarks of PD, alpha-synuclein and Lewy body (LB) deposition, progress from the brainstem and olfactory nuclei through the substantia nigra to the neocortical structures (Braak staging). (10) Spatial transcriptomic studies in post-mortem brains have identified that genes involved in dopamine biosynthesis and microglial function such as *SNCA* and *SCARB2* are differentially expressed in the Braak stage associated regions in the normal and well as PD brains. (7) Regional expression patterns were disrupted in the PD brain, including in areas known to be affected in preclinical PD. (7) Changes were not limited to the dopaminergic neurons; within the microglial transcriptome for instance, specific variants (e.g. *P2RY12*) may be associated with transition to a PD state. (9) Single-cell RNA sequencing of the PD midbrain identified dysfunctional dopaminergic neurons characterized by *CADPS2* overexpression and *TH* under expression, along with glial activation. (8) Notwithstanding these important gains in knowledge gleaned from post-mortem brains, this approach is limited by access to tissue, changes subsequent to post-mortem interval and inability to study longitudinal outcomes. (11), (12) Additionally, presence of multiple cell types in bulk brain tissue confounds the interpretation of gene expression data. (13) Brain biopsies may provide restricted access to the superficial cortex; however, they are not routinely performed in PD. (14) Indeed, the core pathology of PD is dominated by changes in the deeply located basal ganglia structures, which are seldom amenable to biopsy in life. In this context, deep brain stimulation (DBS) provides a window of opportunity to access in vivo tissue from living human brains, and specifically from the subthalamic nucleus (STN). (15)

The STN-DBS procedure involves placing stimulating electrodes into the bilateral STN under stereotactic guidance, during an awake craniotomy. (16) To confirm the placement of electrodes into the dorsolateral motor territory of the STN, two steps are performed during surgery-1) microelectrode recording (MER) to record the typical STN potentials and 2) macrostimulation and monitoring for benefits and adverse effects. (17), (18) The micro and macroelectrodes that have traversed the STN may be a novel source of brain tissue from PD brains, without the need for additional invasive procedures. Early work from two separate groups demonstrated the feasibility of this approach to study regional transcriptomic profiles in vivo. (19), (20)

In this paper, we report the development of an optimised protocol to study differential gene expression using total RNA extracted from DBS microelectrodes that have traversed the basal ganglia structures in post-mortem brains. In life, this technique provides a unique opportunity to probe the regional molecular signatures of PD phenotypes, in combination with longitudinal clinical data and imaging.

## Methods

We developed and validated a protocol for RNA extraction from DBS microelectrodes using three unfixed post-mortem human brains at the Department of Forensic Medicine and Toxicology, All India Institute of Medical Sciences (AIIMS), New Delhi. The study was approved by the Institutional Ethics Committee, AIIMS, New Delhi (IEC No: 15/03.02.2023), and written informed consent was obtained from the next of kin of all participants.

### Tissue Collection

To simulate intraoperative tissue acquisition, a trained functional neurosurgeon introduced combined microrecording and macrostimulating DBS microelectrodes (microTargeting™, FHC Inc., ME, USA; tip diameter 25–150 µm, recording area 50 µm², impedance 0.5–1 MΩ,

To simulate intraoperative tissue acquisition, a trained functional neurosurgeon introduced combined microrecording and macrostimulating DBS microelectrodes (microTargeting™, FHC Inc., ME, USA; tip diameter 25–150 µm, recording area 50 µm², impedance 0.5–1 MΩ, outer diameter 0.55 mm) through the frontal cortex to a depth of approximately 10–15 cm in the direction of the basal ganglia and STN, mimicking the standard DBS surgical trajectory. No image guidance was used, as the aim was methodological optimisation rather than anatomical targeting. Critically, microelectrodes were inserted without a guide tube. In clinical DBS surgery, a rigid guide tube extends from the cortical surface to within a few millimetres of the target, shielding the electrode from contact with traversed tissue so that adherent material derives predominantly from the target region. In this autopsy model, the absence of a guide tube meant the electrode was exposed along its full insertion length, and tissue from frontal cortex, white matter, and all traversed subcortical structures may have adhered to the electrode surface-an important distinction when interpreting transcriptomic profiles. Following a five-minute dwell time, electrodes were withdrawn and immediately transferred into lysis buffer (Buffer RLT with DTT). Several collection strategies were evaluated: (i) recording tip only, (ii) combined recording and stimulating segments, and (iii) electrode washings; and both single and pooled electrode collections. Samples were transferred on ice, centrifuged, and stored at −80°C until processing. **(Figure 1)**

**Figure 1.**
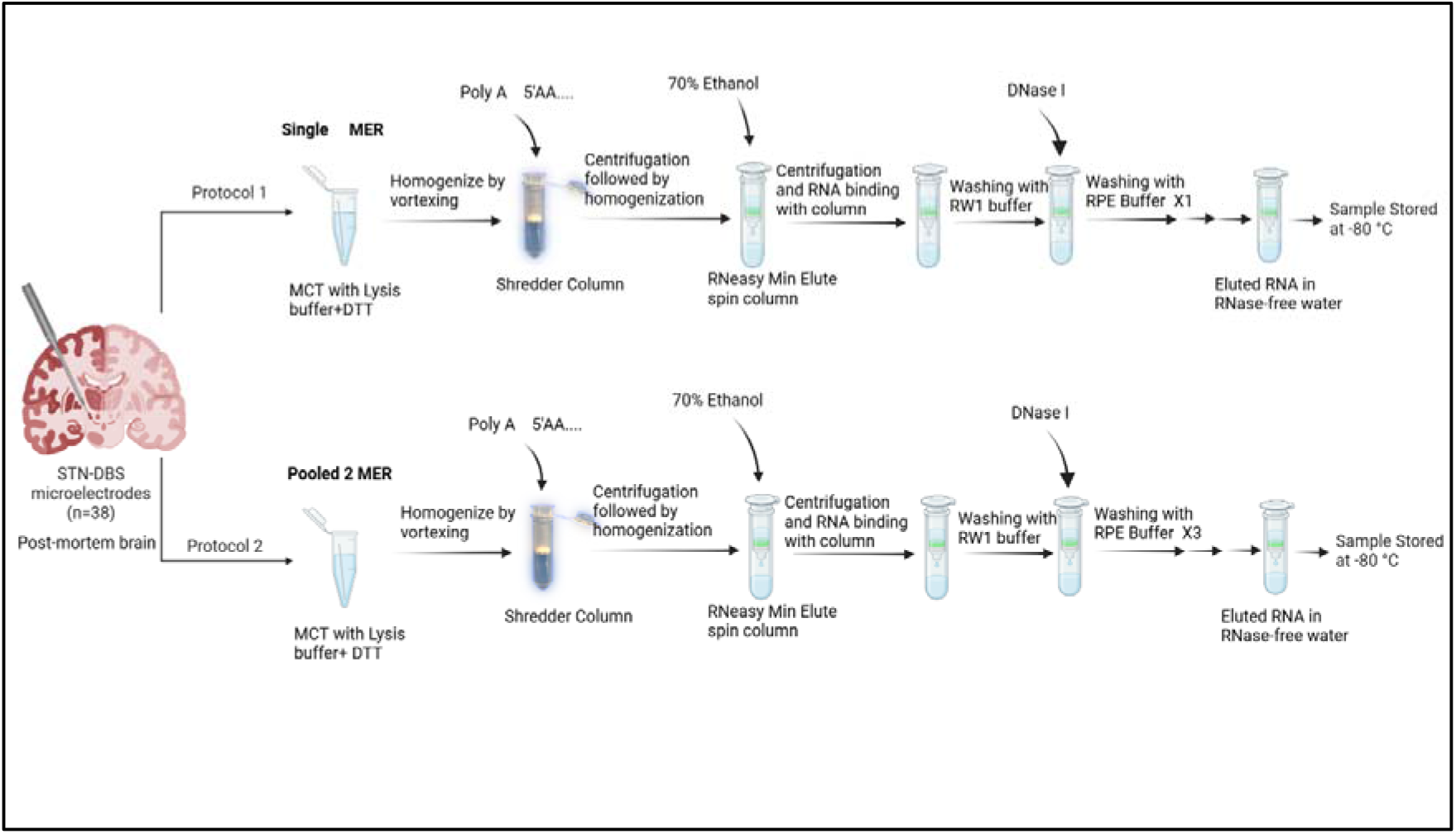
RNA isolation protocol used for high throughput transcriptomics from deep human brains using DBS microelectrodes

### RNA Isolation and Quality Assessment

Total RNA was extracted using a modified RNeasy Micro protocol (Qiagen, Germany). Poly(A) carrier RNA was added to the lysate prior to extraction to enhance recovery. After homogenisation and QIAshredder column centrifugation, the flow-through was mixed with 70% ethanol, loaded onto a RNeasy Mini column, and washed with RW1 buffer. On-column DNase I digestion was performed (15 min, room temperature) to eliminate genomic DNA. RNA was eluted in RNase-free water and stored at −80°C. Concentration and purity were assessed by NanoDrop spectrophotometry (Thermo Fisher Scientific, USA). A subset of samples (n=11, single-electrode collections) was further evaluated using the Agilent 2100 Bioanalyzer with the RNA 6000 Pico kit (Agilent Technologies, USA) to determine RNA Integrity Number (RIN).

### cDNA Library Preparation and Sequencing

Sequencing libraries were prepared using the NEBNext® Single Cell/Low Input RNA Library Prep Kit for Illumina (New England Biolabs, USA). RNA was reverse-transcribed, amplified by 11 PCR cycles, and 100 pg of amplified cDNA subjected to tagmentation and indexing using NEBNext® Single Index Primer sets. Library concentration and quality were assessed using the Qubit™ 2.0 Fluorometer with the HS DNA Assay kit (Thermo Fisher Scientific). Qualified libraries were pooled and sequenced on the Illumina NovaSeq 6000 platform (paired-end, 2 × 150 bp) targeting approximately 15 million read pairs per library.

### Bioinformatic Analysis

Raw reads were processed on the Galaxy platform 26.1.rc1. (21) Quality assessment was performed using Falco and MultiQC. (22), (23) Adapter and low-quality base trimming were carried out using Cutadapt (minimum Phred score 20, minimum read length 20 bp). (24) Trimmed reads were aligned to the human genome (GRCh38) using STAR with GENCODE v49 annotations. (25), (26) Alignment quality was assessed using MultiQC and RSeQC (gene body coverage, read distribution, duplicate assessment with MarkDuplicates). (27), (28) Gene-level counts were generated using featureCounts. (29) Differential gene expression analysis was performed with DESeq2, with p-values adjusted for multiple testing using the Benjamini–Hochberg false discovery rate (FDR); genes with adjusted p-value <0.05 were considered significantly differentially expressed. (30), (31), (32) Data were visualised using PCA, sample-to-sample distance heatmaps, and gene expression heatmaps. The dimensionality of the rlog-transformed expression data was determined by Horn’s Parallel Analysis (1,000 permutations), retaining PCs whose eigenvalues exceeded the 95th percentile of eigenvalues from randomly permuted data of equivalent dimensions. Sample-to-sample distances were computed as Mahalanobis distances in the retained PC space, which accounts for the covariance structure between principal components. **(Supplementary Figure 1)**

### Tissue Identity and Brain Sub-regional Analysis

To confirm brain tissue origin and characterise sub-regional identity, DESeq2-normalised expression profiles were compared against the GTEx v10 reference dataset spanning 54 human tissues. Tissue-specific marker genes were identified using the τ tissue-specificity index computed on GTEx v10 median TPM values across all 54 tissues. (24) Tau scores range from 0 (ubiquitous expression) to 1 (strictly tissue-specific). Genes were selected as markers if they met three criteria: Tau > 0.80, maximum TPM > 5.0 across all tissues, and peak expression in a brain or blood tissue (excluding non-neural high-Tau genes). This yielded 996 marker genes: 768 with peak expression in brain sub-regions and 228 with peak expression in whole blood. Pearson correlation coefficients were computed between log - transformed GTEx marker profiles and log (normalised counts + 0.5) for each autopsy sample. Statistical significance was assessed by permutation testing (1,000 permutations, gene labels shuffled on the GTEx reference profile) with Benjamini-Hochberg FDR correction across all tissue-sample comparisons.

To further characterise sub-regional tissue identity at greater anatomical resolution, normalised expression profiles were compared against the Allen Human Brain Atlas (ABA) normalised microarray dataset (donors H0351.2001 and H0351.2002), which uniquely includes the STN alongside 18 adjacent discrete subcortical regions. (33) A τ-based marker selection approach was not applied to the ABA dataset for two reasons: (i) the ABA covers only brain regions, so τ computed across these similar sub-regions was substantially lower, with a threshold of 0.80 leaving fewer than 40 named genes; and (ii) empirical testing confirmed that at any threshold sufficient to exclude non-specific genes (Tau > 0.50), the ABA yields too few STN-specific markers for stable Pearson correlation. This reflects the fundamental transcriptomic similarity of adjacent subcortical nuclei (mean pairwise r = 0.965 between subcortical and cortical regions) rather than a methodological limitation. A genome-wide correlation approach using all 16,590 genes common to both datasets was therefore used. Pearson correlations were computed between ABA region profiles (log-normalised microarray intensities) and log (normalised counts + 1) for each autopsy sample. Statistical significance was assessed by permutation testing (1,000 permutations, gene labels shuffled on the ABA profile) with Benjamini–Hochberg FDR correction across all region–sample comparisons (n = 406 tests), using df = 13. To quantify transcriptional overlap between subcortical and cortical brain regions, pairwise Pearson correlations were computed between mean ABA profiles of 11 STN-adjacent subcortical and 10 sensorimotor cortical regions, and subcortical-enriched gene filtering was applied to assess whether removing the shared transcriptional background improved regional discrimination.

### Cell-type Deconvolution

To characterise the cellular composition of RNA recovered from DBS microelectrodes, MuSiC (Multi-subject Single-cell deconvolution) (26) was applied using the Allen Human Brain Atlas Human-Mammalian Brain Atlas Basal Ganglia (HMBA-BG) snRNA-seq dataset as a reference. (34) Thirteen cell types from 10 human donors were included: STR D1 MSN, STR D2 MSN, STR Hybrid MSN (medium spiny neurons), M Dopa (midbrain dopaminergic neurons), F Glut and F M Glut (frontal and frontal-midbrain glutamatergic cortical neurons), CN ST18 GABA and STR SST GABA (GABAergic interneurons), Astrocyte, Oligodendrocyte, OPC (Oligodendrocyte Precursor Cells), Microglia, and Endothelial cells. A representative random sample of 500 cells per cell type per donor was drawn from the HMBA-BG reference (total n = 500 cells per group × 10 donors × 13 cell types; random seed 42). The top 5,000 most variable genes common to both the bulk and single-cell datasets (n = 5,000 of 36,601 ABA genes) were used for correlation. Negative proportion estimates, which arise as a MuSiC artefact for near-absent cell types, were set to zero and proportions renormalised to sum to one. Analysis was performed in R (v4.6.0) using MuSiC v1.0.0 loaded via pkgload, with a SingleCellExperiment reference object built from the pseudo-bulk matrix.

## Results

Three post-mortem brains from male donors aged 24–35 years were included (**Supplementary Table 1**). The mean post-mortem interval was 21.3 ± 20.6 days. A total of 38 microelectrodes were inserted across the three brains. In Autopsy Brain 1 (AB1), tissue from 12 individual microelectrodes was processed separately. In AB2 and AB3, pairs of microelectrodes from the right and left hemispheres were pooled prior to extraction, yielding 8 and 5 pooled samples respectively. In total, RNA extraction was performed on 25 samples (12 single, 13 pooled) **(Figure 2).**

**Figure 2:**
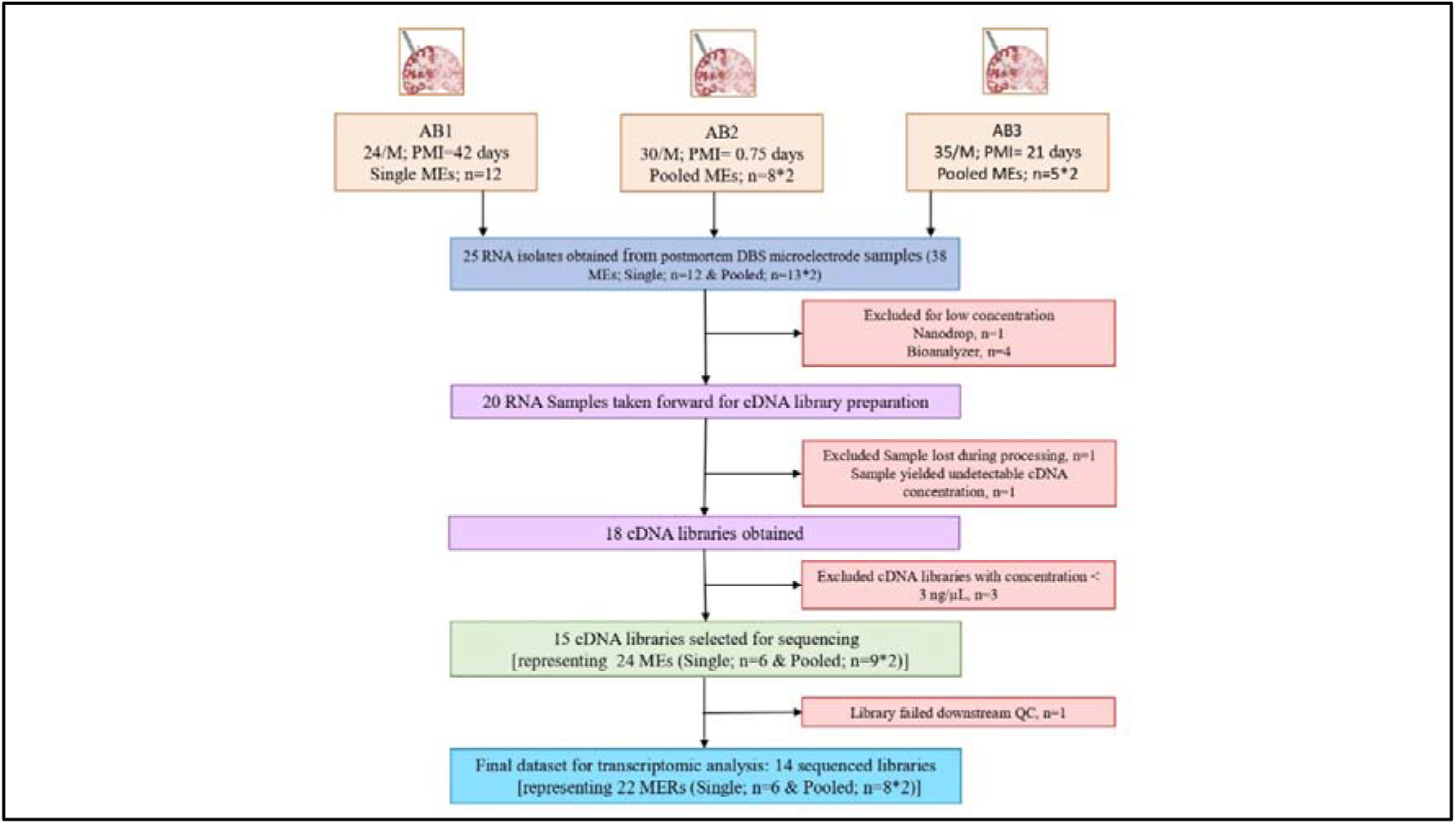
Samples flow chart.

### RNA Yield, Integrity, and Library Quality

All 25 RNA isolates were assessed by NanoDrop spectrophotometry (**Supplementary Table 2**). Mean RNA concentration (n=24) was 4.2 ± 3.1 ng/µL and median A260/A280 ratio was 1.9 (range 0.3–3.8). Eleven single-electrode samples were further assessed by Bioanalyzer; 6/11 (54.5%) achieved RIN ≥5 with a median RIN of 7.1 (range 5.9 - 7.8) (**Supplementary Table 3**). Twenty samples were taken forward for cDNA library preparation; one was lost during processing and one yielded undetectable cDNA, resulting in 18 libraries. Three libraries had concentrations below 3 ng/µL. After quality assessment, 15 libraries representing 24 microelectrodes (6 single, 9 pooled) were selected for sequencing with mean concentration of 39.6 ± 20.6 ng/µL (**Supplementary Table 4**). Mean sequencing depth was 43.0 ± 14.1 million read pairs (range: 11.2–64.2 million); mean Q30 score was 83.2 ± 5.6 % and mean GC content 35.6 ± 4.9 %. One library failed downstream QC and was excluded, leaving 14 libraries for all subsequent analyses.

### Sequencing Quality Control

Gene body coverage analysis (RSeQC) across all 14 libraries showed broadly consistent read distribution from 5′ to 3′ ends, with the majority of samples demonstrating overlapping coverage profiles and no evidence of severe transcript-end enrichment that would indicate widespread RNA degradation (**Figure 3A**). A moderate 3′ bias was observed in a subset of samples, expected given the low-input nature of the material and variable post-mortem RNA integrity. PCA showed partial clustering corresponding to the three autopsy donors, with PC1 accounting for 55.3% and PC2 for 11.8% of total variance (**Figure 3B**). AB1 samples showed the greatest spread along PC1, AB2 and AB3 samples showed distinct yet partly overlapping distributions. Hierarchical clustering of pairwise sample-to-sample distances confirmed intra-donor similarity and inter-donor separation, with AB1 showing the greatest transcriptional distance from AB2 and AB3 (**Figure 3C**). Dispersion modelling showed the expected inverse relationship between dispersion and mean expression, confirming that the DESeq2 statistical framework is well-calibrated for this dataset (**Figure 3D**).

**Figure 3A.**
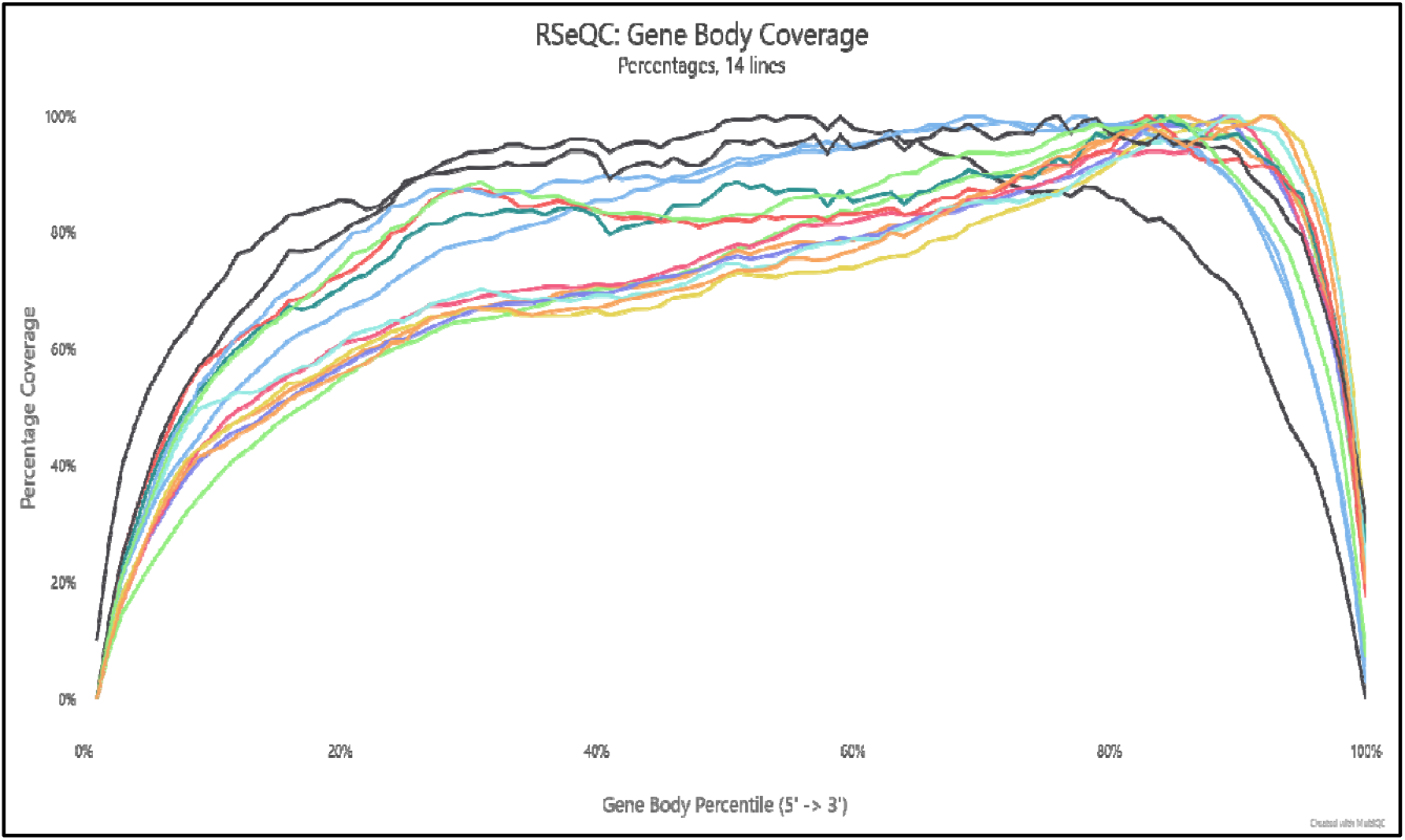
Gene body coverage across 14 samples

**Figure 3B.**
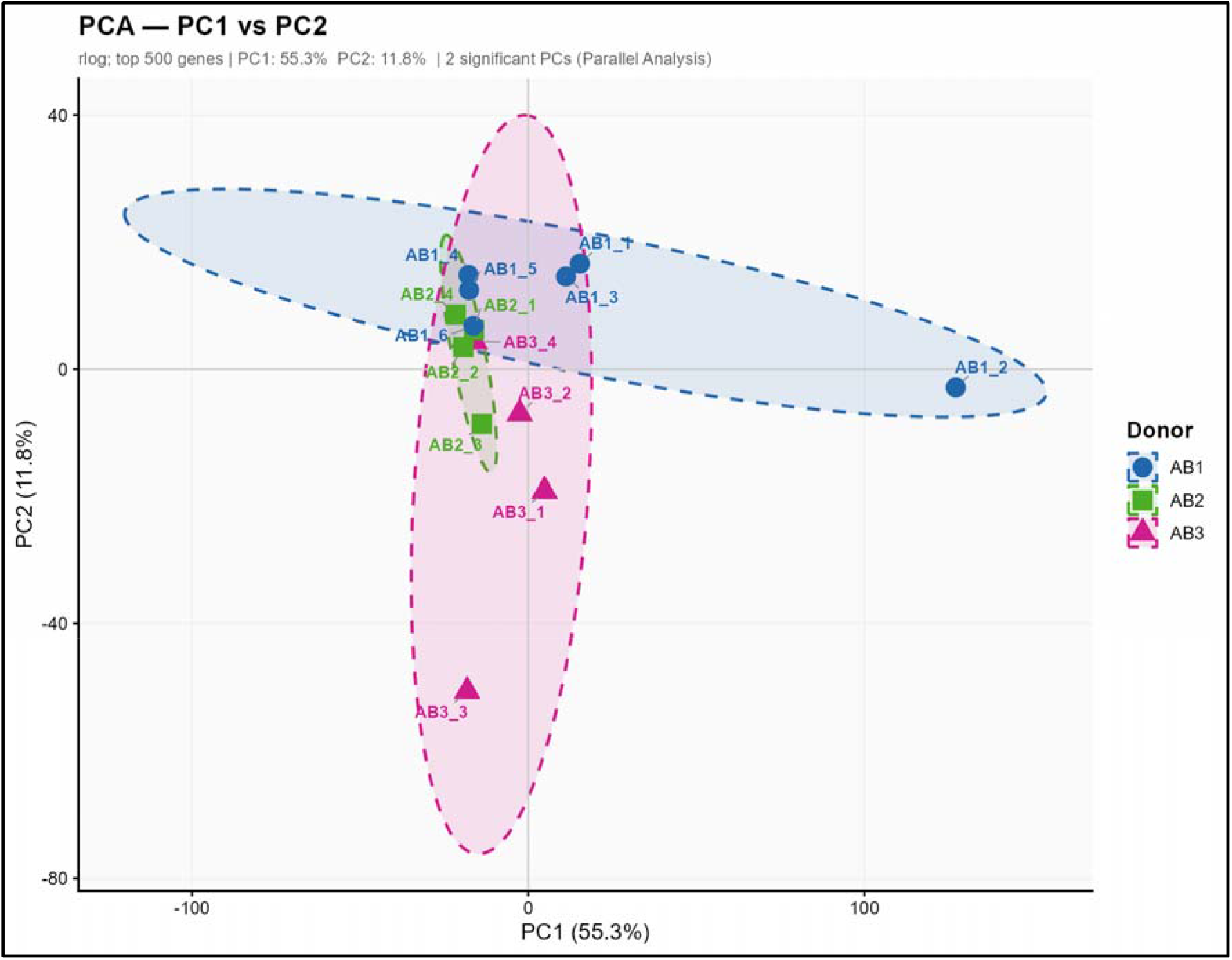
Principal Component Analysis of transcriptomic variation. PCA plot showing separation of samples into AB1 (blue circles), AB2 (green squares), and AB3 (pink triangles) autopsy brain donors. PC1: 55.3%, PC2: 11.8% of variance explained. rlog transformed gene expression counts; top 500 most variable genes.

**Figure 3C.**
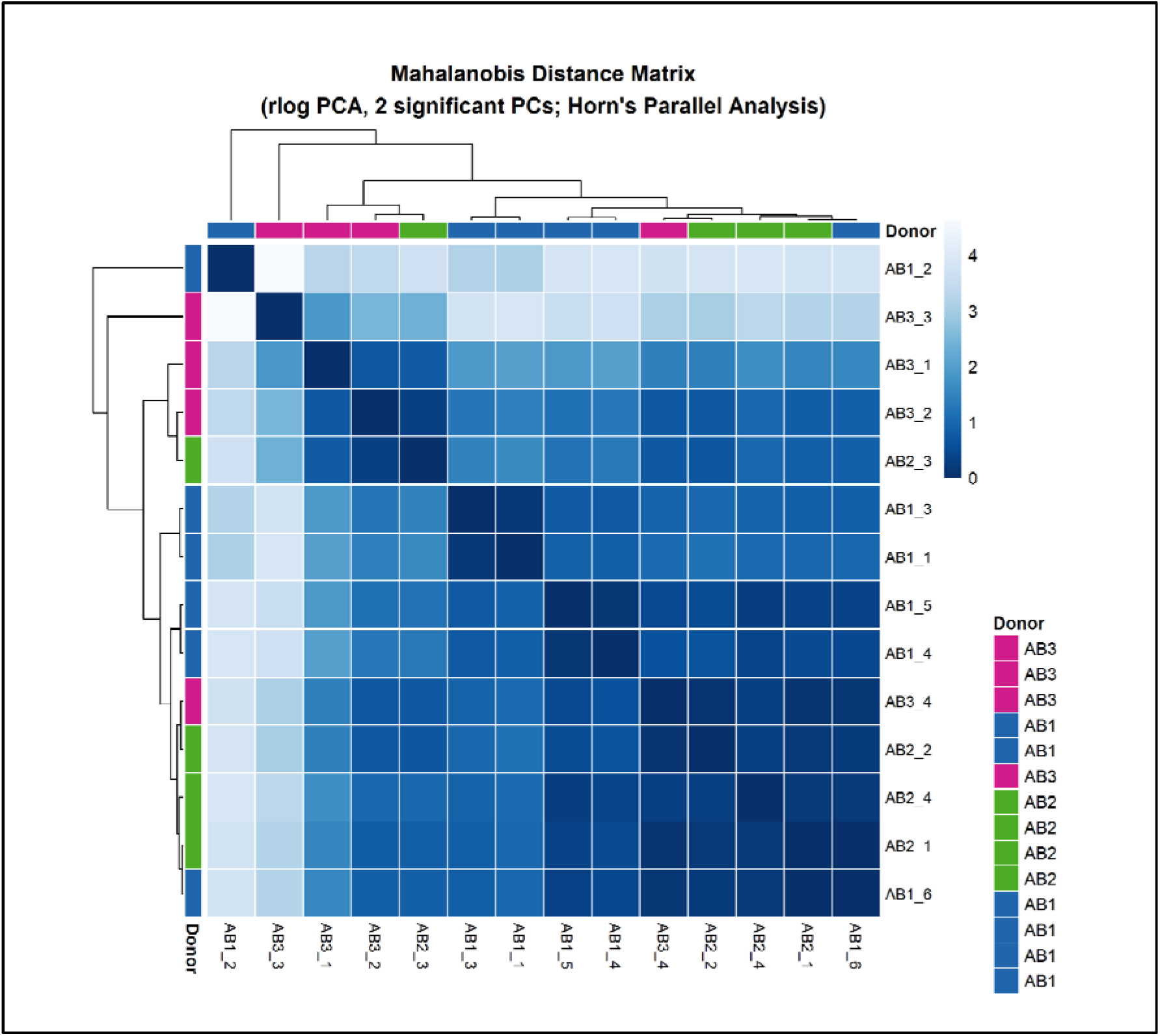
Sample-to-sample distance heatmap. Hierarchical clustering demonstrating high intra-group similarity and moderate inter-group separation.

**Figure 3D:**
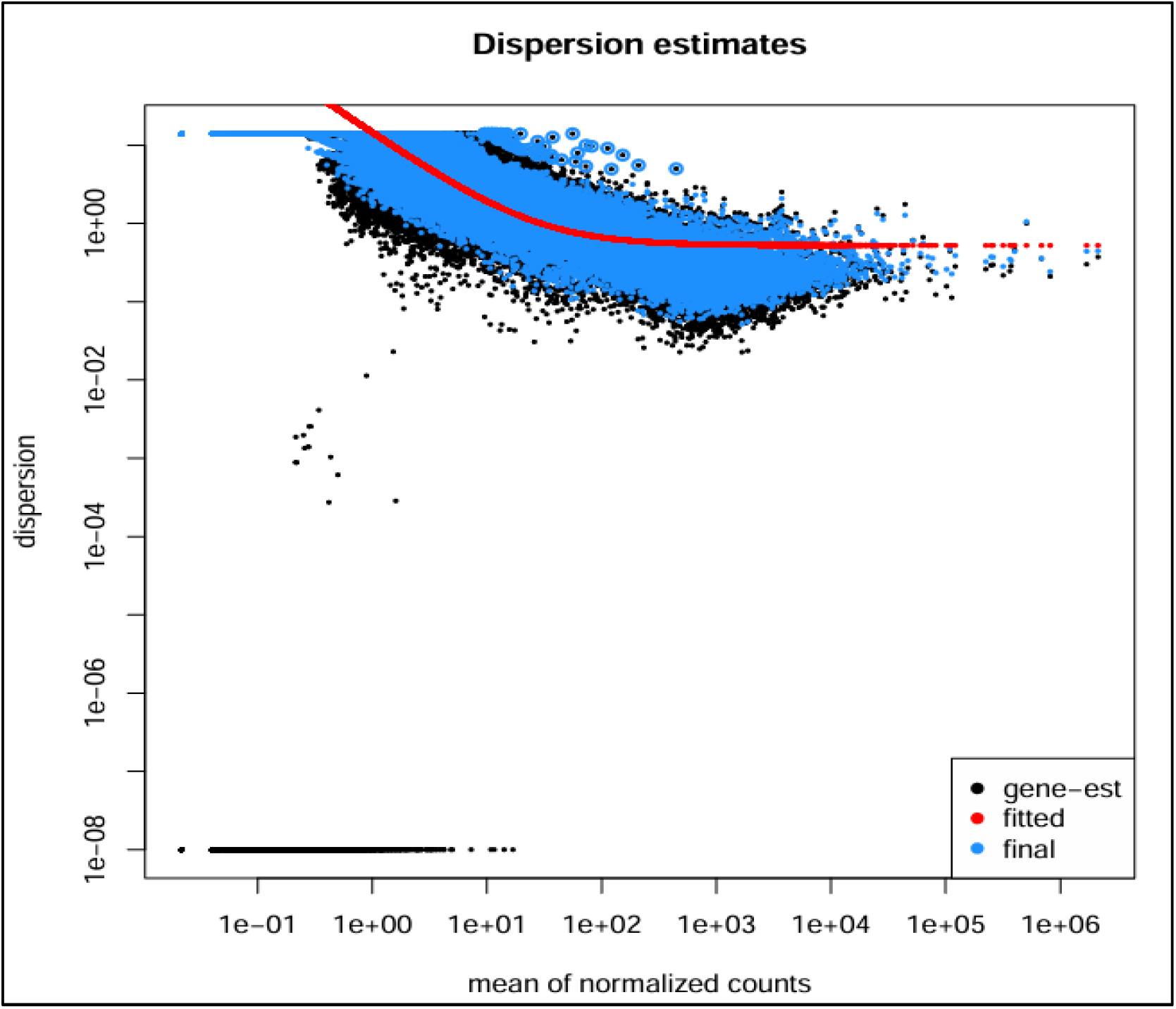
Dispersion estimates for RNA-seq data. Gene-wise and fitted dispersion values across mean expression levels.

### Transcriptomic profiling confirms brain-specific gene expression

To confirm that RNA recovered from DBS microelectrodes was of brain rather than blood or peripheral tissue origin, we assessed the genome-wide expression profiles of all 14 samples against a reference panel of human tissue types from the GTEx dataset (v10), encompassing whole blood, solid organs, and 13 discrete brain sub-regions. The most highly expressed genes by mean normalised count were canonical neuronal and glial markers: *MBP* (baseMean 487573.7 ± 641989.1), *PLP1* (107254.1 ± 72653.21), *SNAP25* (71992.3 ± 31022.4), *NTRK2* (14736. ± 6331.8), *GFAP* (24826.0 ± 17987.3), and *FOXP1* (12006.9 ± 11476.4), all characteristically enriched in brain tissue. Mitochondrial transcripts (MT-*RNR2* (2118149 ± 2493879), MT-*CO2* (1690762 ± 1935836), MT-*ND4* (801264.6 ± 657228.1)) were also among the most abundant, consistent with the high metabolic demands of neural tissue. Hierarchical clustering of TPM-normalised expression values across a curated set of tissue-informative genes revealed that the autopsy brain samples clustered most closely with brain-derived reference tissues, including multiple cortical and subcortical regions, and were clearly separated from non-neural tissues such as whole blood, liver, lung, skin, and reproductive tissues. (**Figure 4**).

**Figure 4:**
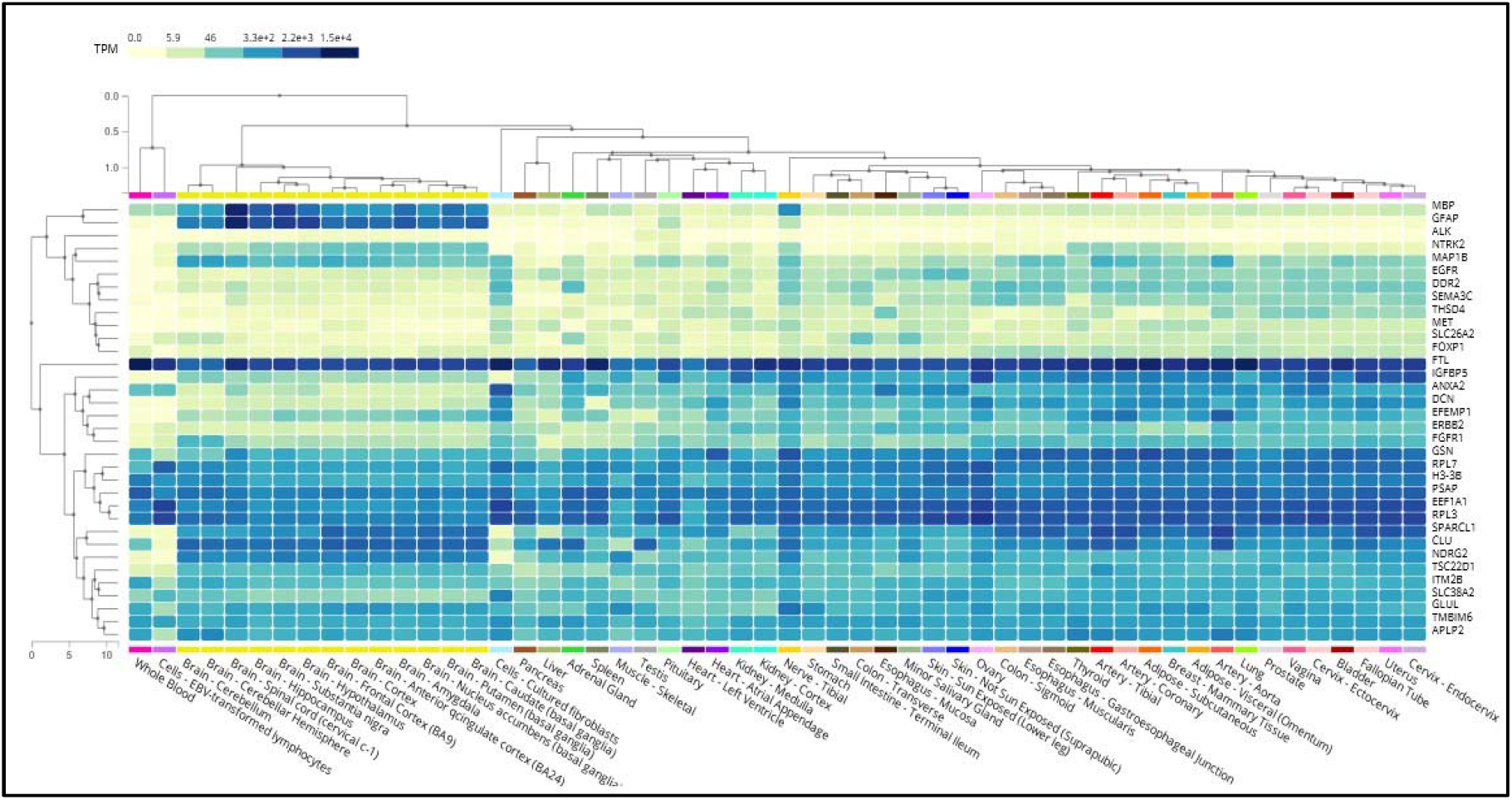
Heatmap of gene expression with cell-type annotation and unsupervised hierarchical clustering across autopsy brain samples

### Brain sub-regional identity

Pearson correlation of each sample’s expression against GTEx tissue profiles revealed a consistent transcriptomic signature across all 14 samples and three donors **(Figure 5)**, all BH-FDR < 0.05). The strongest correlations were with frontal cortex (overall mean r = 0.683 ± 0.072), anterior cingulate cortex (overall mean r = 0.658 ± 0.069), and amygdala (overall mean r = 0.615 ± 0.068). Basal ganglia sub-regions showed consistent positive correlations: caudate (overall mean r = 0.553 ± 0.059), putamen (overall mean r = 0.546 ± 0.059), and nucleus accumbens (overall mean r = 0.544 ± 0.057). Hypothalamus (overall mean r = 0.541 ± 0.050) and substantia nigra (overall mean r = 0.501 ± 0.060) showed moderate positive correlations consistent with STN-adjacent anatomy. Cerebellar profiles showed the lowest brain correlations (cerebellar hemisphere: overall mean r = 0.327 ± 0.039; cerebellum: overall mean r = 0.314 ± 0.037), consistent with the electrode trajectory not reaching the posterior fossa. Strikingly, whole blood correlation was strongly negative across all donors (mean r = −0.250 ± 0.079, BH-FDR < 0.05), providing unambiguous confirmation that the transcriptomic signal does not reflect major haematological contamination.

**Figure 5:**
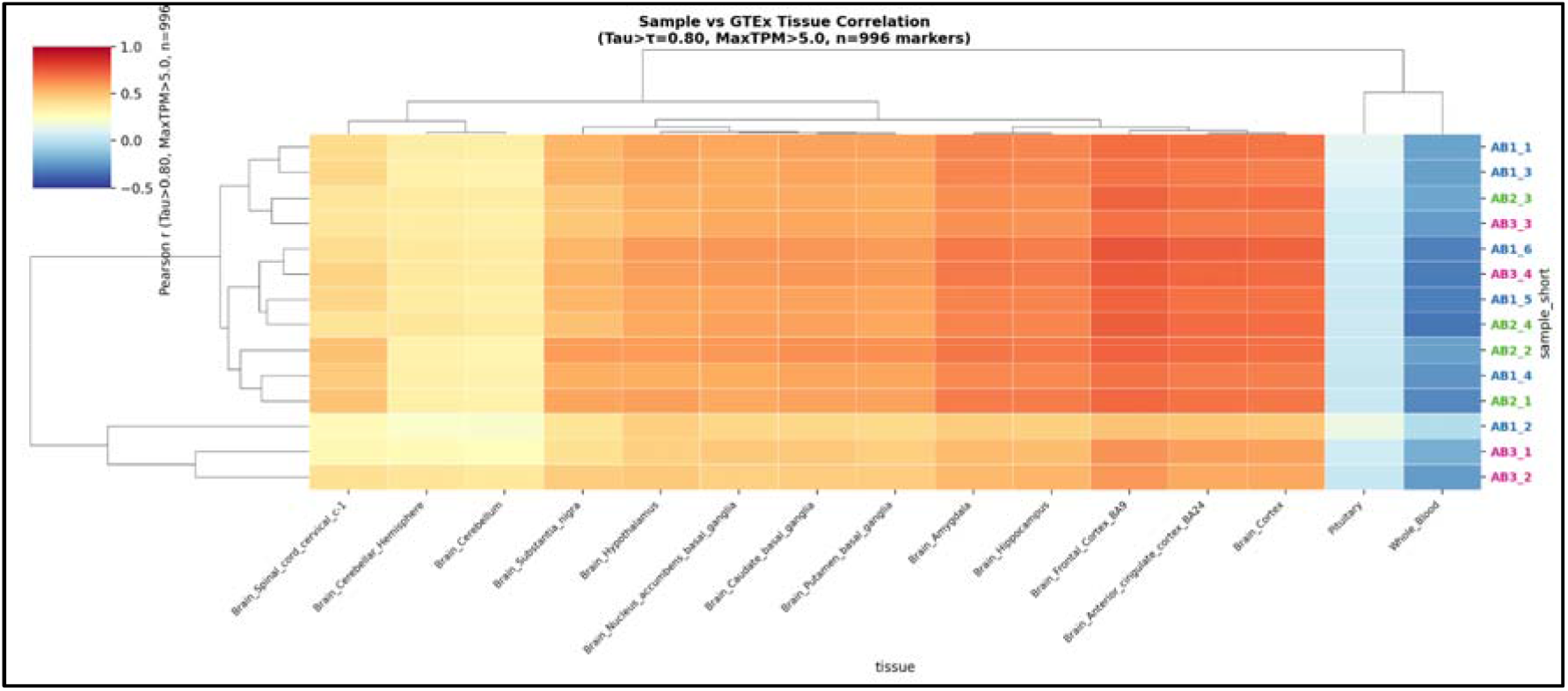
Pearson correlation of each sample’s expression against GTEx tissue profiles, using 996 high-specificity Tau-based marker genes

### Allen Human Brain Atlas sub-regional validation

Genome-wide Pearson correlation against ABA normalised microarray data (16,590 common genes) confirmed positive correlations between autopsy microelectrode samples and all 19 ABA regions, including the STN itself (**Figure 6A, 6B**). All correlations were non-random (permutation p < 0.001 for all region–donor comparisons, BH-FDR < 0.05 for 344/406 per-sample tests). The STN showed consistent correlations across all three donors (mean Pearson r = 0.544 ± 0.127 0.625 ± 0.014, 0.562 ± 0.058 for AB1, AB2, and AB3 respectively, overall mean r=0.572 ± 0.012). Regions with highest overall mean r included nucleus accumbens (0.592 ± 0.059), amygdala (CeA: 0.591 ± 0.059; BLA: 0.589 ± 0.059), globus pallidus external (0.588 ± 0.057), and putamen (0.586 ± 0.058), all immediate anatomical neighbours of the STN. The zona incerta (0.564 ± 0.057) and substantia nigra pars compacta (0.568 ± 0.056)-structures medial and inferior to the STN-also showed strong positive correlations.

**Figure 6.**
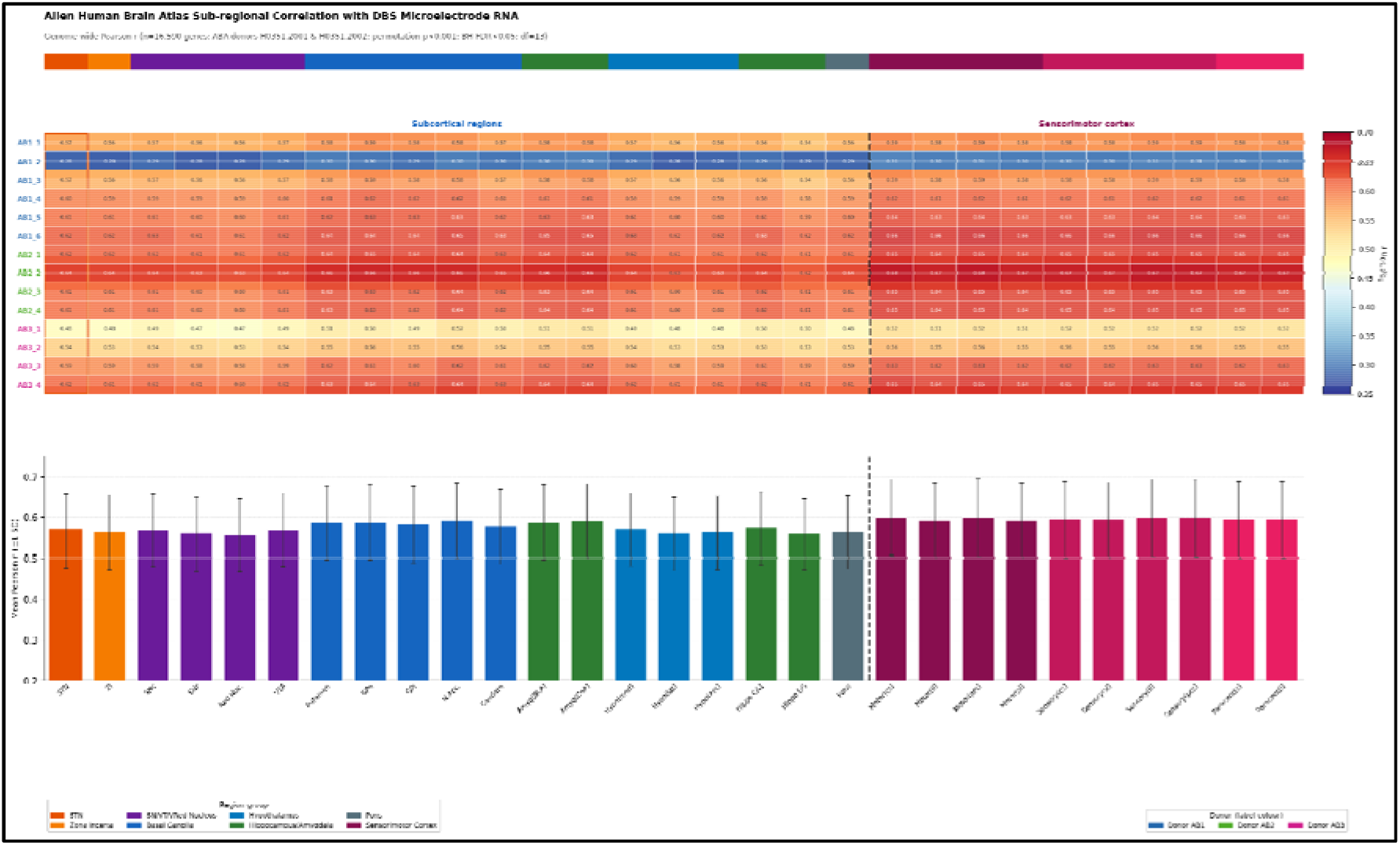
A, 6B: Genome-wide Pearson correlation against Allen Human Brain Atlas normalised microarray data (16,590 common genes)

### Transcriptional overlap between subcortical and cortical regions

To determine whether the narrow spread of ABA correlation values across all regions (0.557– 0.606) reflected genuine tissue mixing or inherent transcriptomic similarity between brain regions, we quantified the genome-wide overlap between subcortical and sensorimotor cortical ABA profiles. Pairwise Pearson correlations between all subcortical-cortical region pairs were strikingly high (mean r = 0.965 ± 0.008, range 0.952–0.982), indicating that subcortical and cortical brain regions share approximately 96% of genome-wide transcriptional variance. Of 29,131 ABA-profiled genes, 20,811 (71.4%) showed no meaningful differential expression between the two groups (|log FC| ≤0.5), with 5,657 genes (19.4%) subcortically enriched and 2,663 (9.2%) cortically enriched. Sensorimotor cortical regions ranked among the highest overall ABA correlations with autopsy samples (mean r = 0.598–0.606), reflecting both this shared transcriptional background and the physical traversal of frontal cortex by the unguided electrode.

To test whether removing the overlapping gene signal would sharpen discrimination, ABA-autopsy correlations were recomputed using only the 5657 subcortical-enriched genes. This did not improve regional discrimination: mean subcortical r = 0.383, cortical r = 0.397, a gap of +0.013 in favour of cortex compared with-0.023 in the genome-wide analysis. The top subcortical-enriched genes included *DDC*, *RGS9*, *HTR2C*, *PBX3*, and *IRX2* - established markers of basal ganglia and diencephalon-while cortical-enriched genes included *THEMIS*, *SLC17A7*, *CAMK2A*, and *NRGN*. The persistence of cortical signal after subcortical enrichment filtering confirms that the mixed transcriptomic profile reflects genuine cortical tissue contribution along the unguided electrode trajectory, rather than non-specific background. In clinical DBS practice, the guide tube would be expected to substantially limit this cortical contribution.

### Cell-type deconvolution of DBS microelectrode RNA

MuSiC cell-type deconvolution against the HMBA-BG snRNA-seq reference revealed a consistent and biologically coherent cellular composition across all 14 samples (**Figure 7A, 7B, 7C**). Oligodendrocytes were the predominant estimated cell type (mean proportion 33.5 ± 17.2%), consistent with the abundant myelinated white matter traversed by the unguided electrode trajectory. STR D1 MSNs were consistently detected across all three donors (mean 6.0 ± 4.9%), confirming the presence of genuine striatal neuronal signal in the recovered RNA. STR D2 MSNs (mean 0.3 ± 0.7%) and STR Hybrid MSNs (mean 1.8 ± 3.2%) were also detected, with AB2 showing the highest striatal MSN proportions across all three subtypes, consistent with the greater striatal transcriptomic signal observed in that donor in the ABA correlation analysis. Midbrain dopaminergic neurons (M Dopa) were present at low but detectable proportions (mean 1.1 ± 2.7%), consistent with the sparse distribution of these cells relative to the primary electrode target. Frontal glutamatergic cortical neurons (F Glut; mean 9.1 ± 7.0%) were detected across all samples, independently corroborating at the single-cell level the transcriptomic evidence of cortical tissue accumulation along the unguided electrode. AB3 showed the highest cortical neuron proportions (F Glut mean 14.5 ± 9.5%), consistent with its greater cortical transcriptomic signature in the GTEx and ABA analyses. Endothelial cells contributed unexpectedly high proportions in several samples (mean 12.1 ± 10.9%), likely reflecting vascular tissue adherent to the electrode surface at the cortical entry point. Glial cell types including astrocytes (mean 6.4 ± 5.2%) and microglia (mean 8.7 ± 5.1%) were consistently present at proportions appropriate for bulk subcortical tissue. The inter-donor pattern of cellular composition closely mirrored the donor-level clustering observed in PCA and hierarchical clustering, confirming that biological differences in anatomical sampling along the electrode trajectory are a primary driver of inter-donor transcriptional variability.

**Figure 7:**
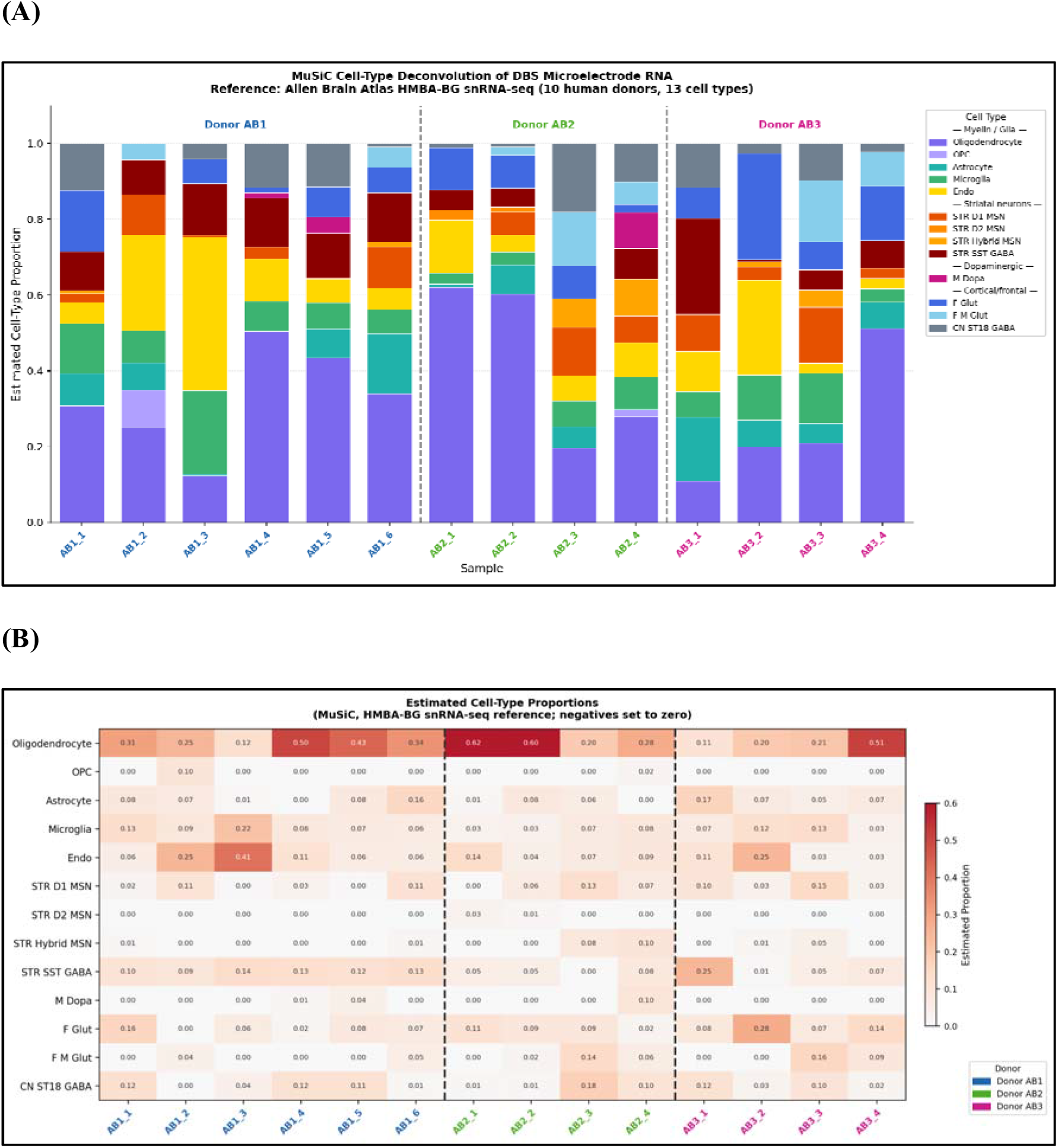
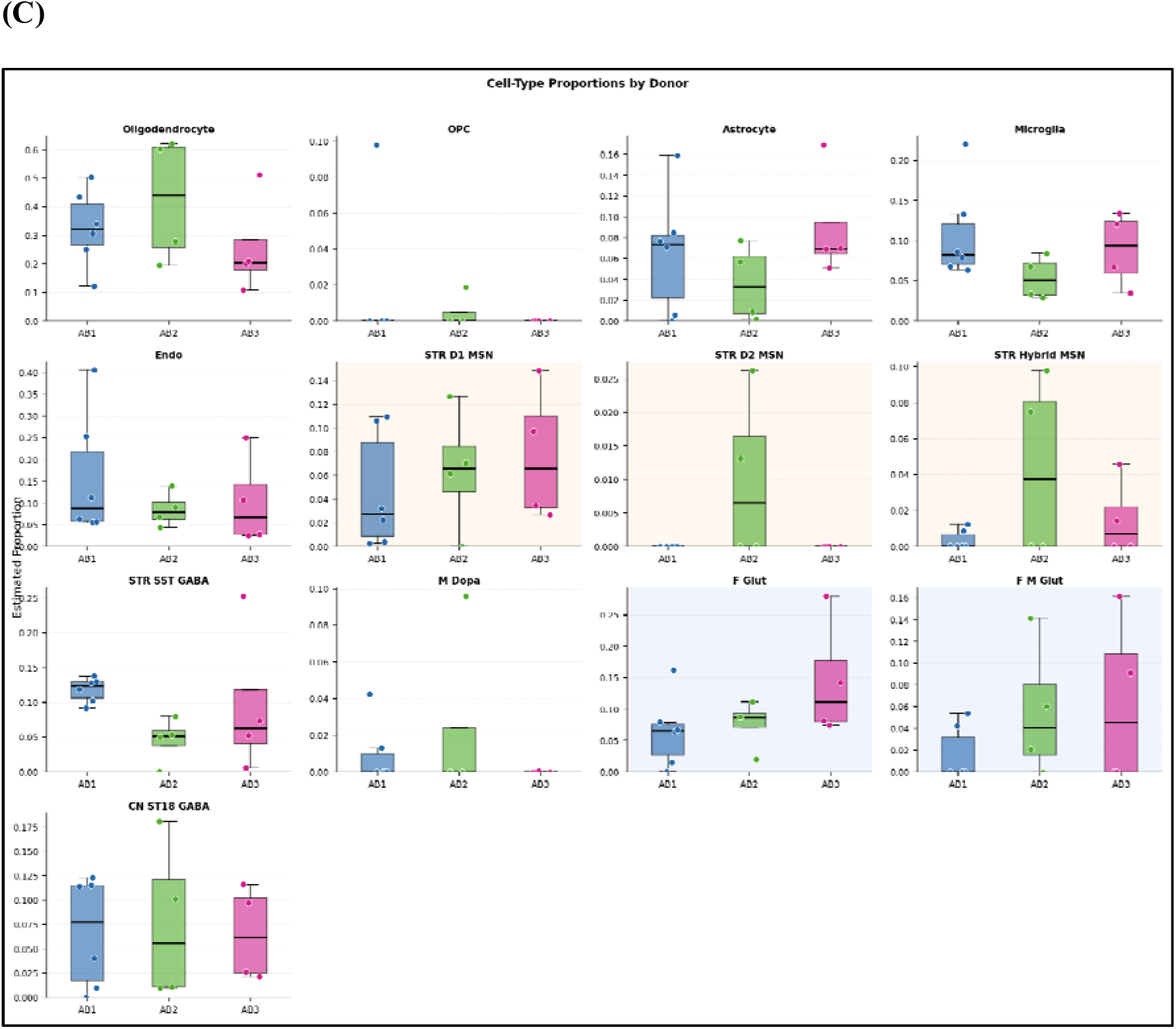
Cell-type deconvolution of DBS microelectrode bulk RNA-seq using MuSiC. **(A)** Estimated cell-type proportions per sample. Stacked bar chart showing MuSiC-estimated proportions of 13 cell types across all 14 autopsy microelectrode samples, grouped by donor (AB1, blue; AB2, green; AB3, pink). Reference single-nucleus RNA-seq data were derived from the Allen Brain Cell Atlas Human-Mammalian Brain Atlas Basal Ganglia dataset (HMBA-BG; 10 human donors). Cell types are grouped by biological category: myelin/glia (Oligodendrocyte, OPC, Astrocyte, Microglia, Endo), striatal neurons (STR D1 MSN, STR D2 MSN, STR Hybrid MSN, STR SST GABA), dopaminergic (M Dopa), and cortical/frontal neurons (F Glut, F M Glut, CN ST18 GABA). **(B)** Heatmap of estimated cell-type proportions. Colour intensity represents the estimated proportion of each cell type (rows) in each sample (columns), ranging from white (0%) to dark red (≥60%). Sample labels are colour-coded by donor, dashed vertical lines separate donors. Values shown in each cell represent the renormalised estimated proportion. **(C)** Cell-type proportions by donor. Boxplots showing the distribution of estimated proportions for each of the 13 cell types across samples within each donor (AB1, blue; AB2, green; AB3, pink). Individual data points represent single microelectrode samples. Orange shading indicates striatal neuronal cell types; blue shading indicates cortical/frontal neuronal cell types. Boxes show the interquartile range; horizontal line indicates the median.

## Discussion

In this study, we demonstrate that high-quality RNA suitable for transcriptomic profiling can be recovered from minimal tissue adherent to DBS microelectrodes using a modified low-input RNA extraction protocol. Despite the inherent challenges of post-mortem brain tissue, including RNA degradation and restricted input material, our workflow produced consistent RNA recovery, reliable library generation, and high-quality sequencing data, establishing a robust foundation for application during live DBS surgery.

The RNA quality metrics observed here are consistent with prior reports demonstrating that moderate integrity values are common in RNA derived from autopsy brain tissue due to intrinsic post-mortem degradation. (35) Importantly, successful library preparation and sequencing were achieved in the majority of isolates, even among samples with suboptimal RIN values, corroborating emerging evidence that RNA-seq is feasible with low-integrity input material when appropriate library preparation strategies are employed. (19) Gene body coverage analysis demonstrated low 5′/3′ bias and broadly consistent coverage across transcripts in most samples, supporting the integrity of the sequencing dataset. Minor deviations in a subset of samples likely reflect inherent variability in RNA quality arising from differences in post-mortem interval and handling conditions, which are well-recognised sources of heterogeneity in autopsy-derived tissue. (36), (37), (38) The partial inter-donor transcriptomic overlap observed in PCA is consistent with the conservation of the human brain transcriptome across healthy individuals, where regional anatomical identity accounts for substantially more variance than individual biological differences. (35)

A critical question for any DBS microelectrode RNA profiling study is whether the extracted RNA is genuinely derived from brain parenchyma rather than from contaminating blood or other non-neural sources. (19) Our brain tissue identity analysis provides robust evidence addressing this question. Genome-wide expression profiling revealed that the most highly expressed genes across all samples were canonical markers of central nervous system cell types, including *MBP* and *PLP1* (oligodendrocytes), *GFAP* (astrocytes), *SNAP25* and *NTRK2* (neurons), and *FOXP1* (striatal neurons), while haematopoietic transcripts were expressed at low levels. (39), (40), (41), (42), (43), (44), (45) Hierarchical clustering against a broad GTEx reference tissue panel confirmed co-clustering with brain tissues and unambiguous separation from whole blood and peripheral organs. (46) The GTEx Tau-based correlation analysis further confirmed brain-specific transcriptomic signatures across all three donors. The frontal cortex ranking highest followed by the anterior cingulate cortex is anatomically expected, as electrodes traversed frontal cortex without a guide tube in this post-mortem model. (47) Basal ganglia sub-regions (caudate, putamen, nucleus accumbens) showed consistent positive correlations consistent with the electrode trajectory approaching the STN through the basal ganglia. (48), (49) Cerebellum ranked lowest among brain tissues and whole blood showed strongly negative correlations, providing a robust and unambiguous negative control that excluded both posterior fossa and haematological contamination.

The ABA analysis extended the sub-regional inference to 19 discrete subcortical regions including the STN. All correlations were statistically non-random (permutation p < 0.001), confirming genuine brain tissue signal throughout. The STN and its immediate neighbours-zona incerta, substantia nigra, globus pallidus, and putamen-all showed consistent positive correlations across donors. Critically, sensorimotor cortical regions showed the highest ABA correlations of all, reflecting the 96% genome-wide transcriptional overlap between subcortical and cortical brain regions. Subcortical-enriched gene filtering did not resolve this ambiguity, confirming that the mixed cortical–subcortical profile is driven by genuine tissue accumulation along the unguided electrode, rather than by non-specific gene expression.

The MuSiC cell-type deconvolution provides the first single-cell-resolution characterisation of tissue composition from DBS microelectrode RNA, and strongly corroborates the bulk transcriptomic findings. (50) The dominance of oligodendrocytes (33.5 ± 17.2%) confirms that the electrode trajectory traverses predominantly myelinated white matter, as expected for a freehand STN-targeting approach through the frontal lobe. (51), (52) The consistent detection of STR D1 MSNs (mean 6.0 ± 4.9%), across all donors provides cell-type-level validation of genuine striatal signal, complementing the ABA genome-wide correlation evidence. The presence of frontal cortical neurons (F Glut, 9.1 ± 7.0%) is explained by the absence of a guide tube leading to cortical tissue accumulation along the electrode. (53) This convergence between bulk transcriptomic, ABA correlational, and deconvolution evidence substantially strengthens the biological plausibility of the findings. The unexpectedly high endothelial proportion in some samples (up to 40%) may reflect differential vascular adherence depending on the electrode insertion trajectory and proximity to minor vascular structures in this non-image guided insertion. (54), (55) In future intraoperative studies using a guide tube and image guidance, endothelial contamination would be expected to diminish, and the STN-specific neuronal signal-particularly D1 and D2 MSNs relevant to basal ganglia circuit dysfunction in PD-should be proportionally enriched. (56), (57)

There are several clinical implications of this study. First, it provides a robust framework to access deep subcortical tissue in vivo, enabling exploration into molecular drivers of phenomenology and treatment response in PD. Second, in combination with spatial transcriptomic mapping this technique maybe utilized for cellular-level image guidance for DBS programming. Third, it can support longitudinal investigations into neurotransmitter and circuitry changes post-DBS with molecular and pathway characterisation of the electrode trajectory at implantation.

Several limitations of the present study should be acknowledged. First, the use of unfixed post-mortem brain tissue introduces variability in RNA quality that would not be present in samples obtained intraoperatively in vivo. The wide range of post-mortem intervals across donors represents an inherent limitation of autopsy-based validation, though the successful recovery of sequenceable RNA even from samples with extended post-mortem intervals supports the robustness of the protocol. Second, microelectrodes were inserted without image guidance or guide tube allowing tissue accumulation from the full trajectory. Future intraoperative studies would use a guide tube, which would be expected to substantially enrich the subcortical transcriptomic signal. Third, all regional correlations were statistically significant but bulk RNA-seq cannot statistically discriminate between adjacent subcortical regions, given 96% genome-wide transcriptional overlap and the limited sample dimensionality. Fourth, the study included three donors, all male and aged 25–35 years without known neurological disease, and therefore does not capture the transcriptomic changes associated with Parkinson’s disease or the older age range typical of DBS candidates. Validation in intraoperative samples from PD patients, will be required before clinical application. Finally, the RNA input quantities recovered from individual microelectrodes were extremely low, and a proportion of samples required pooling or failed library preparation, highlighting the need for continued optimisation of extraction efficiency for application in clinical settings where material is strictly limited.

Taken together, these findings demonstrate that the low-input RNA extraction protocol successfully recovers brain-specific, anatomically coherent transcriptomic profiles from DBS microelectrode tissue. The capacity to discriminate brain from blood at the transcriptomic level, and to identify sub-regional and cellular signatures consistent with the surgical target site, provides the critical proof-of-concept required before applying this approach to RNA recovered from microelectrodes used in patients undergoing DBS for Parkinson’s disease. Future work will apply this protocol in the operative setting to explore the molecular correlates of PD phenotypic heterogeneity in vivo, in combination with longitudinal clinical and neuroimaging data.

## Supporting information

Supplementary Figure 1: Scree plot showing distribution of principal components for normalized gene expression data.

Supplementary Table 1: Autopsy Brain Details

Supplementary Table 2: Qualitative and quantitative measurements of RNA isolates using nanodrop

Supplementary Table 3: Qualitative and quantitative measurements of RNA isolates using Bioanalyzer

## Acknowledgement

We would like to thank Dr. Saumyaranjan Mallick (for 2100 Agilent Bioanalyzer instrument support), Department of Pathology, AIIMS, New Delhi.

## Authors’ Roles

R.R. conceived and supervised the study, secured funding and oversaw project administration. R.R., S.S., A.S., D.M.R., A.E., D.G., A.D., T.R., A.G., R.B., A.K.S., L.R., D.S., R.K., M.S., K.G., and P.S.C. contributed to data acquisition. S.S., A.S., V.C., S.R., and L.R. developed the methodology. R.R., A.S., U.Y., A.K.S., K.B., and H.H. performed the data analysis. R.R., S.S., and A.S. drafted the original manuscript and revised the manuscript. All authors critically reviewed the manuscript, approved the final version, and agreed to be accountable for all aspects of the work. S.S. and A.S. contributed equally to this work.

## Disclosures

### 1.(a) Conflict of Interest: Authors report no potential conflicts of interest

**(b) Funding Source:** This research was funded by DBT/Wellcome Trust India Alliance under the project entitled “Molecular signatures of Parkinson’s disease in vivo: a deep brain stimulation approach”, (Reference No.: IA/CPHI/22/1/506544).

### 2. Financial Disclosures for the previous 12 months

No relevant financial disclosures.

## Ethical Compliance Statement

⁥ This study was approved by the Institutional Ethics Committee, IEC No: 15/03.02.2023.

⁥ We confirm that we have read the Journal’s position on issues involved in ethical publication and affirm that this work is consistent with those guidelines.

