## Supplementary Figure 1: Scree plot showing distribution of principal components for normalized gene expression data. for "Deep Brain Stimulation Microelectrodes as a Source of Human Subcortical RNA: Validation of a Low-Input Transcriptomic Protocol"

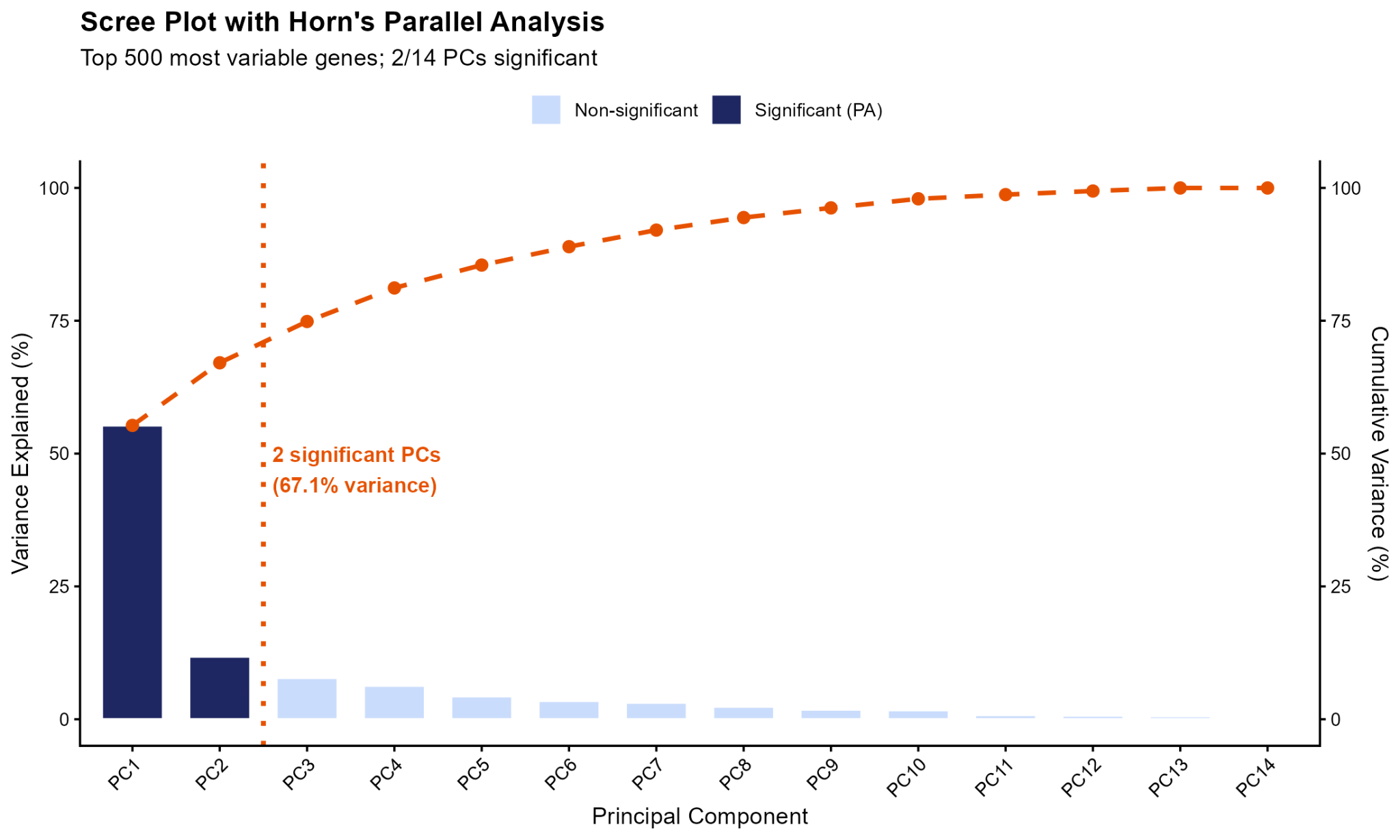

**Supplementary Figure 1:** Scree plot showing distribution of principal components for normalized gene expression data.
