## Supplementary Table 1: Autopsy Brain Details for "Deep Brain Stimulation Microelectrodes as a Source of Human Subcortical RNA: Validation of a Low-Input Transcriptomic Protocol"

| **Subject PM no.** | **Age**  **(Years)** | **Gender** | **PMI** | **Causes of Death** |
| --- | --- | --- | --- | --- |
| AB1 | 20-30 | Male | ~6 Weeks  (42 days) | Sepsis |
| AB2 | 20-30 | Male | ~18 Hours (0.75 days) | Asphyxia |
| AB3 | 30-40 | Male | ~3 Weeks (21 days) | Chronic Lung disease |
