## Supplementary Table 2: Qualitative and quantitative measurements of RNA isolates using nanodrop for "Deep Brain Stimulation Microelectrodes as a Source of Human Subcortical RNA: Validation of a Low-Input Transcriptomic Protocol"

| **S. No.** | **Sample IDs** | **Concentration(ng/µL)** | **A260/280** | **A260/230** | **A260** |
| --- | --- | --- | --- | --- | --- |
| 1 | ADBS001/S01 | 5.430 | 3.818 | 0.012 | 0.1358 |
| 2 | ADBS001/S02 | 4.93 | 1.360 | 0.109 | 0.1232 |
| 3 | ADBS001/S03 | 3.643 | 2.227 | 0.033 | 0.091 |
| 4 | ADBS001/S04 | 3.547 | 1.765 | 0.030 | 0.0887 |
| 5 | ADBS001/S05 | 6.726 | 1.565 | 0.237 | 0.1681 |
| 6 | ADBS001/S06 | 0.427 | 0.305 | 0.0107 | 0.012 |
| 7 | ADBS001/S07 | 5.344 | 1.521 | 0.181 | 0.1336 |
| 8 | ADBS001/S08 | 0.761 | 1.527 | 0.011 | 0.0190 |
| 9 | ADBS001/S09 | 1.230 | 1.740 | 0.071 | 0.0308 |
| 10 | ADBS001/S10 | 0.811 | 3.307 | 0.022 | 0.0203 |
| 11 | ADBS001/S11 | 5.082 | 2.229 | 0.016 | 0.1275 |
| 12 | ADBS001/S12 | 2.424 | 2.206 | 0.031 | 0.0606 |
| 13 | ADBS002/S01 | 4.674 | 2.515 | 0.415 | 0.1169 |
| 14 | ADBS002/S02 | 2.256 | 2.424 | 0.259 | 0.1564 |
| 15 | ADBS002/S03 | 11.816 | 1.624 | 0.548 | 0.2954 |
| 16 | ADBS002/S04 | 3.118 | 0.759 | 0.324 | 0.0754 |
| 17 | ADBS002/S05 | -2.354 | 0.700 | 0.112 | 0.889 |
| 18 | ADBS002/S06 | 1.257 | 1.749 | 0.090 | 0.0314 |
| 19 | ADBS002/S07 | 1.098 | 1.950 | 0.028 | 0.0774 |
| 20 | ADBS002/S08 | 6.734 | 1.981 | 0.408 | 0.1684 |
| 21 | ADBS003/S01 | 3.684 | 2.358 | 0.0921 | 0.0921 |
| 22 | ADBS003/S02 | 4.282 | 3.297 | 0.062 | 0.1071 |
| 23 | ADBS003/S03 | 8.184 | 1.414 | 0.386 | 0.2046 |
| 24 | ADBS003/S04 | 7.488 | 1.753 | 0.316 | 0.1872 |
| 25 | ADBS003/S05 | 6.555 | 1.958 | 0.496 | 0.1639 |
