## Supplementary Table 3: Qualitative and quantitative measurements of RNA isolates using Bioanalyzer for "Deep Brain Stimulation Microelectrodes as a Source of Human Subcortical RNA: Validation of a Low-Input Transcriptomic Protocol"

| **S. No.** | **Sample IDs** | **Concentration(pg/µL)** | **RIN Number** |
| --- | --- | --- | --- |
| 1 | ADBS001/S01 | 203 | 5.8 |
| 2 | ADBS001/S02 | 254 | 6.8 |
| 3 | ADBS001/S03 | 382 | 7.3 |
| 4 | ADBS001/S04 | 111 | 1 |
| 5 | ADBS001/S05 | 49 | NA |
| 6 | ADBS001/S06 | 63 | 2.5 |
| 7 | ADBS001/S07 | 55 | 1 |
| 8 | ADBS001/S08 | 59 | 1 |
| 9 | ADBS001/S09 | 108 | 6.8 |
| 10 | ADBS001/S10 | 110 | 7.8 |
| 11 | ADBS001/S11 | 208 | 7.4 |
