## Supplemental Data 1 for "Deep Brain Stimulation Microelectrodes as a Source of Human Subcortical RNA: Validation of a Low-Input Transcriptomic Protocol"

**Supplementary Table 4:** Quantitative measurements of cDNA libraries using Qubit® 2.0 Fluorometer

| **S. No.** | **Sample IDs** | **Pooled/Single** | **No. of MERs harvested** | **Macro/Micro/Washing** | **cDNA Library conc. (ng/ µL)** |
| --- | --- | --- | --- | --- | --- |
| 1 | ADBS001/S01 | Single | 1 | Macro | 6.38 |
| 2 | ADBS001/S02 | Single | 1 | Macro | 9.32 |
| 3 | ADBS001/S03 | Single | 1 | Macro | 17.4 |
| 4 | ADBS001/S09 | Single | 1 dip only | Washing | 41.5 |
| 5 | ADBS001/S10 | Single | 1 dip only | Washing | 47.0 |
| 6 | ADBS001/S11 | Single | 1 dip only | Washing | 49.6 |
| 7 | ADBS002/S03 | Pooled | 2 | Micro+ Macro | 56.0 |
| 8 | ADBS002/S04 | Pooled | 2 | Micro+ Macro | 59 |
| 9 | ADBS002/S05 | Pooled | 2 | Micro | 38 |
| 10 | ADBS002/S06 | Pooled | 2 | Micro | 39.2 |
| 11 | ADBS002/S07 | Pooled | 2 | Micro | 3.26 |
| 12 | ADBS003/S02 | Pooled | 2 | Micro+ Macro | 50.4 |
| 13 | ADBS003/S03 | Pooled | 2 | Micro+ Macro | 54.7 |
| 14 | ADBS003/S04 | Pooled | 2 | Micro+ Macro | 58 |
| 15 | ADBS003/S05 | Pooled | 2 | Micro+ Macro | 64.5 |
